# A high density linkage map reveals sexual dimorphism in recombination landscapes in red deer (*Cervus elaphus*)

**DOI:** 10.1101/100131

**Authors:** Susan E. Johnston, Jisca Huisman, Philip A. Ellis, Josephine M. Pemberton

## Abstract

High density linkage maps are an important tool to gain insight into the genetic architecture of traits of evolutionary and economic interest, and provide a resource to characterise variation in recombination landscapes. Here, we used information from the cattle genome and the 50K Cervine Illumina BeadChip to inform and refine a high density linkage map in a wild population of red deer (*Cervus elaphus*). We constructed a predicted linkage map of 38,038 SNPs and a skeleton map of 10,835 SNPs across 34 linkage groups. We identified several chromosomal rearrangements in the deer lineage relative to sheep and cattle, including six chromosome fissions, one fusion and two large inversions. Otherwise, our findings showed strong concordance with map orders in the cattle genome. The sex-averaged linkage map length was 2739.7cM and the genome-wide autosomal recombination rate was 1.04cM per Mb. The female autosomal map length was 1.21 longer than that of males (2767.4cM vs 2280.8cM, respectively). Sex differences in map length were driven by high female recombination rates in peri-centromeric regions, a pattern that is unusual relative to other mammal species. This effect was more pronounced in fission chromosomes that would have had to produce new centromeres. We propose two hypotheses to explain this effect: (1) that this mechanism may have evolved to counteract centromeric drive associated with meiotic asymmetry in oocyte production; and/or (2) that sequence and structural characteristics suppressing recombination in close proximity to the centromere may not have yet evolved at neo-centromeres. Our study provides insight into how recombination landscapes vary and evolve in mammals, and will provide a valuable resource for studies of evolution, genetic improvement and population management in red deer and related species.

**Article Summary:** We present a high density linkage map (>38,000 markers) in a wild population of Red deer ***(Cervus elaphus).*** Our investigation of the recombination landscape showed a marked difference in recombination rates between the sexes in proximity to the centromere, with females showing an unusually elevated rate relative to other mammal species. This effect is most pronounced in chromosomes that would have produced a new centromere in the deer lineage. We propose that the observed effects have evolved to counteract selfish genetic elements associated with asymmetrical female meiosis.

## Introduction

The advent of affordable next-generation sequencing and SNP-typing assays allows large numbers of polymorphic genetic markers to be characterised in almost any system. A common challenge is how to organise these genetic variants into a coherent order for downstream analyses, as many approaches rely on marker order information to gain insight into genetic architectures and evolutionary processes (Ellegren, 2014). Linkage maps are often an early step in this process, using information on recombination fractions between markers to group and order them on their respective chromosomes (Sturtevant, 1913; Lander and Schork, 1994). Ordered markers have numerous applications, including: trait mapping through quantitative trait locus (QTL) mapping, genome-wide association studies (GWAS) and regional heritability analysis (Bérénos *et al.,* 2015; Fountain *et al.,* 2016); genome-scans for signatures of selection and population divergence (Bradbury *et al.,* 2013; McKinney *et al.,* 2016); quantification of genomic inbreeding through runs of homozygosity (Kardos *et al.,* 2016); and comparative genomics and genome evolution (Brieuc *et al*., 2014; Leitwein *et al.,* 2016). Linkage maps also provide an important resource in *de novo* genome assembly, as they provide information for anchoring sequence scaffolds and allow prediction of gene locations relative to better annotated species (Fierst, 2015).

One application of high density linkage maps is the investigation of variation in contemporary recombination landscapes. Meiotic recombination is essential for proper disjunction in many species (Hassold and Hunt, 2001; Fledel-Alon *et al.,* 2009); it also generates new allelic combinations upon which selection can act, and prevents the accumulation of deleterious mutations (Muller, 1964; Felsenstein, 1974; Charlesworth and Barton, 1996). Linkage maps have shown that recombination rates can vary within and between chromosomes, populations and species in a wide variety of taxa (Stapley *et al.,* 2008; Kawakami *et al.,* 2014; Smukowski and Noor, 2011). One striking observation is that sex is consistently one of the strongest correlates with recombination rate and landscape variation. The direction and degree of sex differences in recombination, known as “heterochiasmy”, can differ over relatively short evolutionary timescales, and whilst broad trends have been observed (e.g. increased recombination in females), many exceptions remain (Lenormand and Dutheil, 2005; Brandvain and Coop, 2012). Theoretical explanations for the evolution of heterochiasmy include haploid selection, sex-specific selection and sperm competition (Lenormand and Dutheil, 2005; Trivers, 1988; Burt, 2000), but empirical support for each of these theories had been limited (Mank, 2009). One emerging hypothesis is the role of meiotic drive, where asymmetry in cell division during oogenesis can be exploited by selfish genetic elements (i.e. variants which enhance their own transmission relative to the rest of the genome) associated with centromere “strength” (Brandvain and Coop, 2012). Strong centromeres have increased levels of kinetochore proteins, and will preferentially be drawn to one pole of the oocyte, which will become an egg or a polar body, resulting in biased transmission at the stronger/weaker centromere, respectively (Pardo-Manuel De Villena and Sapienza, 2001; Chmátal *et al.,* 2014). Theoretical work has shown that higher female recombination at centromeric regions may counteract drive by increasing the uncertainty at which linked genomic regions segregate into the egg (Haig and Grafen, 1991). As linkage map data for non-model species continues to proliferate, it is now increasingly possible to investigate the key hypotheses for recombination rate variation and heterochiasmy in a wider variety of taxa.

Nevertheless, creating linkage maps of many thousands of genome-wide markers *de novo* is a computationally intensive process requiring pedigree information, sufficient marker densities over all chromosomes and billions of locus comparisons. Furthermore, the ability to create a high resolution map is limited by the number of meioses in the dataset; as marker densities increase, more individuals are required to resolve genetic distances between closely linked loci (Kawakami *et al.,* 2014). Whilst *de novo* linkage map assembly with large numbers of SNPs is possible (Rastas *et al.,* 2016), one approach to ameliorate the computational cost and map resolution is to use genome sequence data from related species to inform initial marker orders. Larger and finer scale rearrangements can then be refined through further investigation of recombination fractions between markers.

In this study, we use this approach to construct a high density linkage map in a wild population of red deer (*Cervus elaphus*). The red deer is a large deer species widely distributed across the northern hemisphere, and is a model system for sexual selection and behaviour (Clutton-Brock *et al.,* 1982; Kruuk *et al.,* 2002), hybridisation (Senn mid Pemberton, 2009), inbreeding (Huisman *et al.,* 2016) and population management (Frantz *et al.,* 2006). They are also an increasingly important economic species farmed for venison, antler velvet products and trophy hunting (Brauning *et al.,* 2015). A medium density map (~600 markers) is available for this species, constructed using microsatellite, RFLP and allozyme markers in a red deer × Père David’s deer (*Elaphurus davidianus*) F_2_ cross (Slate *et al.,* 2002). However, these markers have been largely superseded by the development of a Cervine Illumina BeadChip which characterises 50K SNPs throughout the genome (Brauning *et al*., 2015). SNP positions were initially assigned relative to the cattle genome, but the precise order of SNPs in red deer remains unknown. Here, we integrate pedigree and SNP data from a long-term study of wild red deer on the island of Rum, Scotland to construct a predicted linkage map of ~38,000 SNP markers and a “skeleton” linkage map of ~11,000 SNP markers that had been separated by at least one meiotic crossover. As well as identifying strong concordance with the cattle genome and several chromosomal rearrangements, we also present evidence of strong female-biased recombination rates at peri-centromeric regions of the genome which is more pronounced in fission chromosomes that would have had to produced new centromeres. We discuss the implications of our findings for other linkage mapping studies, and the potential drivers of recombination rate variation and sexual dimorphism of this trait within this system.

## Materials and Methods

### Study Population and SNP dataset

The red deer population is located in the North Block of the Isle of Rum, Scotland (57°02‘N, 6°20‘W) and has been subject to an on-going individual-based study since 1971 (Clutton-Brock *et al.,* 1982). Research was conducted following approval of the University of Edinburgh’s Animal Welfare and Ethical Review Body and under appropriate UK Home Office licenses. DNA was extracted from neonatal ear punches, post-mortem tissue, and cast antlers (see Huisman *et al.,* 2016 for full details). DNA samples from 2880 individuals were genotyped at 50,541 SNP loci on the Cervine Illumina BeadChip (Brauning *et al.,* 2015) using an Illumina genotyping platform (Illumina Inc., San Diego, CA, USA). SNP genotypes were scored using Illumina GenomeStudio software, and quality control was carried out using the *check.marker* function in GenABEL v1.8-0 (Aulchenko *et al.,* 2007) in R v3.3.2, with the following thresholds: SNP geno-typing success >0.99, SNP minor allele frequency >0.01, and ID genotyping success >0.99. A total of 38,541 SNPs and 2,631 IDs were retained. The function identified 126 pseudoautosomal SNPs on the X chromosome (i.e. markers showing autosomal inheritance patterns). Any heterozygous genotypes at non-pseudoautosomal X-linked SNPs within males were scored as missing. A pedigree of 4,515 individuals has been constructed using microsatellite and SNP data using the software Sequoia (Huisman, 2017; see Huisman *et al.,* 2016 for information on deer pedigree construction).

### Linkage map construction

A standardised sub-pedigree approach was used for linkage map construction (Johnston *et al.,* 2016). The pedigree was split as follows: for each link between a focal individual (FID) and an offspring, a sub-pedigree was constructed that included the FID, its parents, the offspring, and the other parent of the offspring (Figure 1), and were retained where all five individuals were SNP genotyped. This pedigree structure characterises crossovers occurring in the gamete transferred from the FID to that offspring. In cases where an individual had more than one offspring, an individual pedigree was constructed for each FID – offspring relationship. A total of 1355 sub-pedigrees were constructed, allowing characterisation of crossovers in gametes transmitted to 488 offspring from 83 unique males and 867 offspring from 259 unique females. Linkage mapping was conducted using an iterative approach using the software CRI-MAP v2.504a (Green *et al.,* 1990), with input and output processing carried out using the R package crimaptoolsv0.1 (S.E.J., available https://github.com/susjoh/crimaptools) implemented in R v3.3.2. In all cases, marker order was specified in CRI-MAP based on the criteria outlined in each section below. In order to ensure that sex-differences in map lengths are not due to the over-representation of female meioses in the dataset, maps were reconstructed for ten subsets of 483 male and 483 female FID-offspring pairs randomly sampled with replacement from the dataset.

**Fig. 1.**
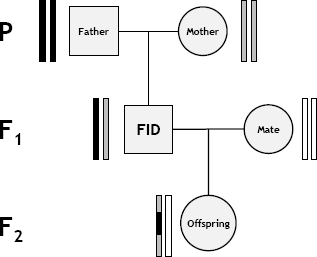
Sub-pedigree structure used to construct linkage maps. Rectangle pairs next to each individual represent chromatids, with black and grey shading indicating chromosome or chromosome sections of FID paternal and FID maternal origin, respectively. White shading indicates chromatids for which the origin of SNPs cannot be determined. Crossovers in the gamete transferred from the focal individual (FID) to its offspring (indicated by the grey arrow) can be distinguished at the points where origin of alleles origin flips from FID paternal to FID maternal and vice versa. From Johnston *et al*. (2016).

#### Build 1: Order deer SNPs based on synteny with cattle genome

Mendelian incompatibilities were identified using the CRI-MAP *prepare* function, and incompatible genotypes were removed from both parents and offspring. SNPs with Mendelian error rates of >0.01 were discarded (N = 0 SNPs). Sub-pedigrees with more than 50 Mendelian errors between an FID and its offspring were also discarded (N = 4). All SNPs were named based on direct synteny with the cattle genome (BTA vUMD 3.0; N = 30). Therefore, loci were ordered and assigned to linkage groups assuming the cattle order, and a sex-averaged map of each chromosome was constructed using the CRI-MAP *chrompic* function (N = 38,261 SNPs, Figure S1).

#### Build 2: Rerun cattle order with wrongly positioned chunks removed

All SNP loci from Build 1 were assigned to “chunks”, defined as a run of SNPs flanked by map distances of ≥3 centiMorgans (cM). Several short chunks were flanked by large map distances, indicating that they were wrongly positioned in Build 1 (Figure S1); chunks containing <20 SNPs were removed from the analysis for Build 2 (N = 327 SNPs). A sex-averaged map of each chromosome was reconstructed using the *chrompic* function (N = 37,934 SNPs, Figure S2).

#### Build 3: Arrange chunks into deer linkage groups

SNPs from Build 2 were arranged into 34 deer linkage groups (hereafter prefixed with CEL) based on a previous characterisation of fissions and fusions from the red deer × Père David’s deer linkage map (Slate *et al.,* 2002) and visual inspection of linkage disequilibrium (LD, R^2^, calculated using the *r2fast* function in GenABEL; Figure S3). At this stage, the orientation of linkage groups was made to match that of the Slate et al. publication. There was strong conformity with fissions and fusions identified in the previous deer map (Table 1); intra-marker distances of ~100 cM between long chunks indicated that they segregated as independent chromosomes. In Build 2, chunks flanked by gaps of ≪100cM but >10cM were observed on the maps associated with BTA13 (CEL23) and BTA28 (CEL15; Figure S2). Visual inspection of LD indicated that these chunks were incorrectly orientated segments of ~10.5 and ~24.9 cM in length, respectively (Figure S3a and S3b; Table 1). Reversal of marker orders in these regions resulted in map length reductions of 19.4 cM and 20.9 cM, respectively. Visual inspection of LD also confirmed fission of CEL19 and CEL31 (syntenic to BTA1), with a 45.4cM inversion on CEL19 (Figure S3c). The X chromosome (BTA30, CEL34) in Build 2 was more fragmented, comprising 9 large chunks (Figure S4). Visual inspection of LD in females indicated that chunks 3 and 7 occurred at the end of the chromosome, and that chunks 4, 5 and 6 were wrongly-oriented (Figure S5). After rearrangement into new marker orders, a sex-averaged map of each deer linkage group was reconstructed using the *chrompic* function (N = 37,932 SNPs, Figure S6).

**Table 1:** Synteny between the cattle and deer genomes. Large-scale fissions and fusions are informed by Slate et al. 2002 and confirmed in this study through sequence alignment (Table S5).y Indicates where fission chromosomes would have had to have formed a new centromere.

| Deer Linkage Group (CEL) | Cattle Chr (BTA) | Sheep Chr (OAR) | Notes |
| --- | --- | --- | --- |
| 1 | 15 | 15 |  |
| 2 | 29 | 21 |  |
| 3 | 5 | 3* | Fission from CEL22 in deer lineage. |
| 4 | 18 | 14 |  |
| 5 | 17, 19 | 17, 11 | Fusion of BTA17 (OAR17) & BTA19 (OAR11) in deer lineage. Likely to be the metacentric chromosome in deer. |
| 6 | 6 | 6 | Fission from CEL17 in deer lineage. † |
| 7 | 23 | 20 |  |
| 8 | 2 | 2* | Fission from CEL33 in deer lineage. † |
| 9 | 7 | 5 |  |
| 10 | 25 | 24 |  |
| 11 | 11 | 3* |  |
| 12 | 10 | 7 |  |
| 13 | 21 | 18 |  |
| 14 | 16 | 12 |  |
| 15 | 26, 28 | 22, 25 | Fission into BTA26 (OAR22) & BTA28 (OAR25) in the early cattle/sheep lineage (Slate *et al.*, 2002). On segment syntenic with BTA28, ~13Mb inversion in deer lineage and ~1.5Mb inversion in cattle lineage. |
| 16 | 8 | 2* | Fission from CEL29 in deer lineage. † |
| 17 | 6 | 6 | Fission from CEL6 in deer lineage. |
| 18 | 4 | 4 |  |
| 19 | 1 | 1* | Fission from CEL31 in deer lineage, followed by ~36Mb inversion. † |
| 20 | 3 | 1* |  |
| 21 | 14 | 9 |  |
| 22 | 5 | 3 | Fission from CEL3 in deer lineage. † |
| 23 | 13 | 13 | ~5.9Mb inversion in cattle lineage. |
| 24 | 22 | 19 |  |
| 25 | 20 | 16 |  |
| 26 | 9 | 8 | Fission from CEL28 in deer lineage. † |
| 27 | 24 | 23 |  |
| 28 | 9 | 8, 9* | Fission from CEL26 in deer lineage. |
| 29 | 8 | 2* | Fission from CEL16 in deer lineage. |
| 30 | 12 | 10 |  |
| 31 | 1 | 1* | Fission from CEL19 in deer lineage. |
| 32 | 27 | 26 |  |
| 33 | 2 | 2* | Fission from CEL8 in deer lineage. |
| 34 (X) | X | X | Three possible translocations (two in deer, one in cattle) and one possible ~18Mb inversion in cattle lineage; see Figure S5. |

\* Sheep chromosomes OAR1, OAR2 and OAR3 are fusions of BTA1 & BTA3, BTA2 & BTA8, BTA5 & BTA11, respectively. OAR9 has a translocation from its homologue of BTA9 to its homologue of BTA14.

#### Build 4: Solve minor local re-arrangements

Runs of SNPs from Build 3 were re-assigned to new chunks flanked by map distances of ≥1 cM. Maps were reconstructed to test whether inverting chunks of <50 SNPs in length and/or the deletion of chunks of <10 SNPs in length led to decreases in map lengths by ≥1cM. One wrongly-orientated chunk of 25 SNPs was identified on CEL15 (homologous to part of the inversion site identified on BTA28 in Build 3), and the marker order was amended accordingly (reducing the map length from 101.4 cM to 98.1 cM). Three chunks on the X chromosome (CEL34) shortened the map by ≥1cM when inverted and were also amended accordingly, reducing the X-chromosome map by 10.8 cM relative to Build 3. The deletion of 35 individual SNPs on 14 linkage groups shortened their respective linkage maps by between 1cM and 6.3cM. A sex-averaged map of each deer linkage group was reconstructed using the *chrompic* function (N = 37,897 SNPs, Figure S7).

#### Build 5: Determining the location of unmapped markers and resolving phasing 191 errors

In Builds 1 to 4, 372 SNPs in 89 chunks were removed from the analysis. To determine their likely location relative to the Build 5 map, LD was calculated between each unmapped SNP and all other SNPs in the genome to identify its most likely linkage group. The CRI-MAP *chrompic* function provides information on SNP phase (i.e. where the grandparent of origin of the allele could be determined) on chromosomes transmitted from the FID to offspring. The correlation between allelic phase was calculated for each unmapped marker and all markers within a 120 SNP window around its most likely position. A total 186 SNPs in 18 chunks could be unambiguously mapped back to the genome; for all other markers, their most likely location was defined as the range in which the correlation of allelic phase with mapped markers was >0.9 (Adjusted *R^2^*). A provisional sex-averaged map of each deer linkage group was reconstructed using the *chrompic* function (N = 38,083 SNPs). Marker orders were reversed on the deer fission linkage groups 6, 8, 16, 22 and 31 to match the orientation of the cattle genome.

Errors in determining the phase of alleles can lead to incorrect calling of double crossovers (i.e. two or more crossovers occurring on the same chromosome) over short map distances, leading to errors in local marker order. To reduce the likelihood of calling false double crossover events, runs of grandparental origin consisting of a single SNP (resulting in a false double crossover across that SNP) were recoded as missing (Figure S8) and the *chrompic* function was rerun. Of the remaining double crossovers, those occurring over distances of ≤ 10cM (as measured by the distance between markers immediately flanking the double crossover) were also recoded as missing. Our justification is that the majority of crossovers captured should be subject to some degree of crossover interference (i.e. Class I crossovers; Phadnis *et al.,* 2011); for the purposes of creating a broad-scale map, we have removed any crossovers that may not have been subject to interference and inflate map distances in this dataset. Finally, sex-averaged and sex-specific maps of each deer linkage group were reconstructed using the *chrompic* and *map* functions (Figure 2, Figure S9).

**Figure 2:**
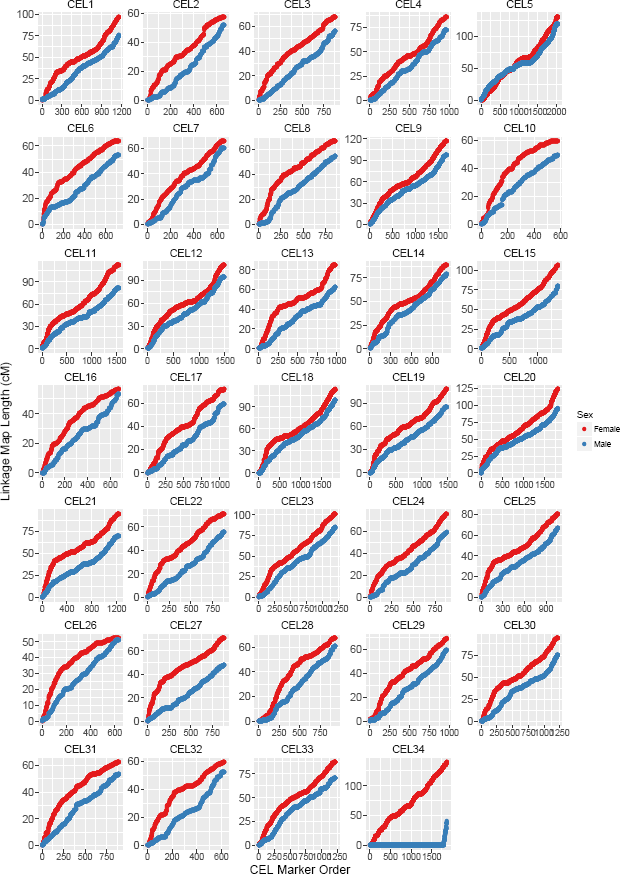
Sex-specific linkage maps for *Cervus elaphus* (CEL) linkage groups after Build 5. Map data is provided in Tables 1, 2 and S1. CEL34 corresponds to the X chromosome; the short map segment in male deer indicates the pseudoautosomal region (PAR).

#### Build 6: Building a skeleton map and testing fine-scale order variations

In Build 5, 71.6% of intra-marker distances were 0cM; therefore, a “skeleton map” was created to examine local changes in marker orders. All runs of SNPs were re-assigned to new chunks where all SNPs mapped to the same cM position; of each chunk, the most phase-informative SNP was identified from the *.loc* output from the CRI-MAP *prepare* function (N = 10,835 SNPs). The skeleton map was split into windows of 100 SNPs with an overlap of 50 SNPs, and the CRI-MAP *flips* function was used to test the likelihood of marker order changes of 2 to 5 adjacent SNPs (*flips2* to *flips5*). Rearrangements improving the map likelihood by >2 would have been investigated further; however, no marker rearrangement passed this threshold and so the Build 5 map was assumed to be the most likely map order (Map provided in Table S1).

#### Determining the lineage of origin of chromosome rearrangements

Lineage of origin and/or verification of potential chromosomal rearrangements was attempted by aligning SNP flanking sequences (as obtained from Brauning *et al.,* 2015) to related genome sequences using BLAST v2.4.0+ (Camacho *et al.,* 2009). Cattle and sheep diverged from deer ~ 27.31 Mya, and diverged from each other ~ 24.6 Mya (Hedges *et al.,* 2015); therefore, rearrangements were assumed to have occurred in the lineage that differed from the other two. Alignments were made to cattle genome versions vUMD3.0 and Btau_4.6.1, and to the sheep genome Oar_v3.1 using default parameters in *blastn,* and the top hit was retained where ≥ 85% of bases matched over the informative length of the SNP flanking sequence.

### Variation in recombination rate and landscape

Estimated genomic positions were calculated for each SNP based on the differences between the cattle base pair position of sequential markers. At the boundaries of rearrangements, the base pair difference between markers was estimated assuming that map distances of 1cM were equivalent to 1 megabase (Mb). The first SNP on each linkage group was given the mean start position of all cattle chromosomes. Estimated genomic positions are given in Table S1. The relationship between linkage map and estimated chromosome lengths for each sex were estimated using linear regression in R v3.3.2.

To investigate intra-chromosomal variation in recombination rates, the probability of crossing over was determined within 1 Mb windows using the estimated genomic positions, starting with the first SNP on the chromosome. This was calculated as the sum of recombination fractions *r* within the window; the *r* between the first and last SNPs and each window boundary was calculated as 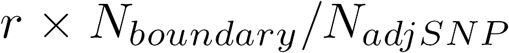, where 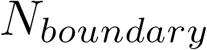 is the number of bases to the window boundary and 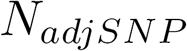 is the number of bases to the adjacent window SNP. Windows with recombination rates in the top 1 percentile after accounting for chromosome size were removed, as very high recombination rates may indicate map misassembly and/or underestimation of physical distances. All deer chromosomes are acrocentric, with the exception of one unknown autosome (Gustavsson and Sundt, 1968). The Build 5 linkage groups maps were orientated in the same direction as the cattle genome, and so we assumed that centromere positions in deer were at the start of the chromosome, as in cattle (Band *et al.,* 2000; Ma *et al.,* 2015).

In fission events, assuming no change in the original centromere position, one fission chromosome would have retained the centromere (in this case, CEL3, 17, 28, 29, 31 and 33), whereas the other would have had to have formed a new centromere (CEL22, 6, 26, 16, 19 and 8, respectively). As all acrocentric chromosomes showed a consistently high recombination rate around the female centromere (see Results), we assumed that neo-centromeres had positioned themselves at the beginning of these chromosomes. We defined chromosome histories as follows: those with fissions retaining the old centromere; fissions that would have formed a new centromere; and chromosomes with no fission or fusion relative to sheep/cattle lineages. Comparison of recombination landscapes between chromosomes of different histories was carried out using general additive models (GAM) from 0Mb (centromere) to 40Mb, specifying k = 10, using the R library mgcv v1.8-15 (Wood, 2011) implemented in R v3.3.2. Recombination rates within each bin were adjusted for chromosome size by dividing the bin rates by the overall chromosome recombination rate (cM/Mb) for each sex. As these chromosome comparisons have a relatively small sample size (n = 32), the GAM analysis was repeated (a) excluding each chromosome and (b) excluding two chromosomes in turn, in order to determine whether the observed effect was driven by one or two chromosomes, respectively. As chromosome sizes are markedly different between fissions retainining a centromere and those forming a new centromere (see Figure S10), comparisons were also made between new centromere chromosomes and unchanged chromosomes of similar size (in this case, CEL6, 8, 16, 22 and 26 vs. CEL2, 7, 10, 24, 27 and 32).

### Transmission distortion

We conducted a preliminary analysis to identify regions of the genome associated with transmission distortion in the red deer pedigree. Specifically, we wished to determine if regions in close proximity to centromeres had biased transmission, which if occurring close to centromeric regions, may indicate differences in centromere strength in the contemporary pedigree. At a given locus, the specific allele transmitted from an FID to a given offspring can be identified in cases where the FID is heterozygous and its mate is homozygous. For each locus per FID sex, an exact binomial test was used to determine whether the transmission frequency of allele A relative to allele B was significantly different from that expected due to chance. The associated P values were transformed to follow an approximate normal distribution using the equation 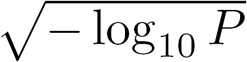, and each SNP locus was assigned to a 1Mb bin. General linear models were run in males and females separately for the 10Mb interval in closest proximity to the centromere on all acrocentric chromosomes, including an interaction term between chromosome history and bin identity.

### Data availability

The Supplemental Material contains information on additional analyses conducted and is referenced within the text. Table S1 contains the full red deer linkage maps for both sexes, including estimated Mb positions and information on marker informativeness. Table S2 contains comparisons of red deer linkage map positions with cattle and sheep genomes for the X chromosome. Table S3 contains the approximate positions of unmapped loci. Table S4 contains the probabilities of crossing over within 1Mb windows in both sexes. Table S5 contains BLAST results to determine lineage of origin of chromosome rearrangements. Table S6 contains the per-locus results for the transmission distortion analysis. Raw data, supplementary tables, and sequence information is publicly archived at doi:10.6084/m9.figshare.5002562. Code for the analysis is archived at https://github.com/susjoh/DeerMapv4.

## Results

### Linkage map

The predicted sex-averaged red deer linkage map contained 38,083 SNP markers over 33 autosomes and the X chromosome (Figure 2; Full map provided in Table S1), and had a sex- averaged length of 2739.7cM (Table 2). A total of 71.6% of intra-marker recombination fractions were zero, and so a skeleton map of 10,835 SNPs separated by at least one meiotic crossover was also characterised (Table S1). The female autosomal map was 1.21 times longer than in males (2767.4 cM and 2280.8 cM, respectively, Table 2). In the autosomes, we observed six chromosomal fissions, one fusion and two large and formerly uncharacterised inversions occurring in the deer lineage (Table 1, Figure S3). Otherwise, the deer map order generally conformed to the cattle map order. The X chromosome had undergone the most differentiation from cattle, with evidence of three translocations, including two in the deer lineage and one in the cattle lineage, and one inversion in the cattle lineage (Figure S5, Table S2), although we cannot rule out that this observation is a result of poor assembly of the cattle genome (Zimin *et al.,* 2009). The estimated positions of 90 unmapped markers are provided in Table S4. The BLAST results for determining lineage of origin are provided in Table S5.

**Table 2:** Marker numbers and sex-averaged and sex-specific map lengths for each deer linkage group in Build 5. The estimated length (Mb) of each linkage group is calculated based on homologous SNP positions on the cattle genome BTA vUMD 3.0 and the sheep genome Oar_v3.1.

| Deer Linkage Group (CEL) | Number of Loci | Estimated Length (Mb) | Sex-averaged map length (cM) | Male map length (cM) | Female map length (cM) |
| --- | --- | --- | --- | --- | --- |
| 1 | 1158 | 82.7 | 88.7 | 75.2 | 96.7 |
| 2 | 663 | 50.3 | 55.4 | 51.6 | 57.5 |
| 3 | 885 | 57.7 | 63.8 | 56.5 | 67.8 |
| 4 | 971 | 65.2 | 81.3 | 72.5 | 85.9 |
| 5 | 2039 | 137.9 | 126.8 | 119.7 | 130.8 |
| 6 | 723 | 52.6 | 59.6 | 52.8 | 63.5 |
| 7 | 660 | 51.7 | 64 | 60.6 | 65.7 |
| 8 | 860 | 58 | 62.1 | 54.4 | 66.7 |
| 9 | 1690 | 111.8 | 109.4 | 96.7 | 116.7 |
| 10 | 580 | 42.7 | 55.3 | 49.1 | 59.2 |
| 11 | 1547 | 107.1 | 101.3 | 81.7 | 112.1 |
| 12 | 1486 | 102.1 | 104.2 | 94 | 110 |
| 13 | 986 | 69.8 | 76.3 | 61.9 | 84.3 |
| 14 | 1113 | 82.2 | 85 | 79.4 | 88.2 |
| 15 | 1357 | 96.4 | 96.4 | 79.2 | 105.9 |
| 16 | 674 | 47 | 54.8 | 52.8 | 56.2 |
| 17 | 1059 | 68.3 | 67 | 59 | 71.5 |
| 18 | 1831 | 120.7 | 108 | 98.8 | 113.3 |
| 19 | 1476 | 101.9 | 99.3 | 85.1 | 107.3 |
| 20 | 1810 | 118.6 | 112.9 | 95.6 | 122.9 |
| 21 | 1236 | 84.1 | 85.5 | 69.7 | 94.6 |
| 22 | 882 | 62.3 | 65.2 | 55.2 | 71.1 |
| 23 | 1200 | 83.3 | 95.1 | 84.6 | 101.1 |
| 24 | 885 | 61.3 | 69.7 | 59.1 | 75.9 |
| 25 | 1066 | 72.1 | 76 | 66.9 | 80.6 |
| 26 | 633 | 41.7 | 51.7 | 50.9 | 52.2 |
| 27 | 886 | 62.5 | 62.2 | 47.8 | 70.7 |
| 28 | 938 | 65.5 | 64.7 | 60.3 | 67.2 |
| 29 | 969 | 67.2 | 65.9 | 59.2 | 69.4 |
| 30 | 1220 | 86.2 | 86.9 | 74.4 | 94 |
| 31 | 892 | 57.7 | 59.1 | 53.3 | 62.3 |
| 32 | 623 | 46.7 | 56.7 | 52.5 | 59.2 |
| 33 | 1220 | 80.4 | 80.8 | 70.3 | 86.9 |
| 34 | 1865 | 148.2 | 148.7 | 40 | 138.9 |
| All | 38083 | 2644.1 | 2739.7 | 2320.8 | 2906.3 |
| All autosomal | 36218 | 2495.7 | 2591.1 | 2280.8 | 2767.4 |

### Variation in recombination rate and landscape

There was a linear relationship between estimated chromosome length and sex-averaged linkage map lengths (Adjusted R^2^ = 0.961, Figure 3A). Smaller chromosomes had higher recombination rates (cM/Mb, Adjusted R^2^ = 0.387, Figure 3B), which is likely to be a result of obligate crossing over. Female linkage maps were consistently longer than male linkage maps across all autosomes (Adjusted R^2^ = 0.907, Figures S11) and correlations between estimated map lengths and linkage map lengths were similar in males and females (Adjusted R^2^ = 0.910 and 0.954, respectively; Figure S12). There was no significant difference between the true and sampled map lengths in males and females (Figure S13), suggesting that the data structure did not introduce bias in estimating sex-specific map lengths.

**Figure 3:**
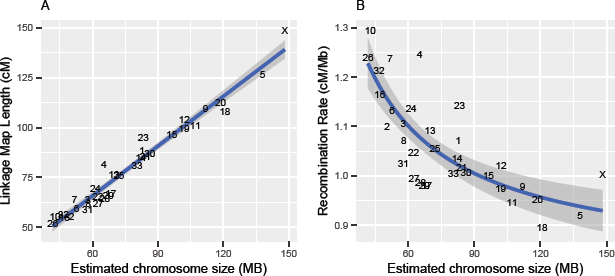
Broad-scale variation in recombination rate, showing correlations be-tween (A) sex-averaged linkage map length (cM) and estimated chromosome length (Mb) and (B) estimated chromosome length (Mb) and chromosomal recombination rate (cM/Mb). Points are chromosome numbers, and lines and the grey-shaded areas indicate the regression slopes and standard errors, respectively.

Fine-scale variation in recombination rate across chromosomes was calculated in 1Mb windows across the genome; recombination rate was considerably higher in females in the first ~20% of the chromosome, where the centromere is likely to be situated (Figure 4). This effect was consistent across nearly all autosomes (Figure 5). Male and female recombination rates were not significantly different across the rest of the chromosome, although male recombination was marginally higher than females in sub-telomeric regions (i.e. where the centromere was absent; Figure 4). Both sexes showed reduced recombination in sub-telomeric regions - this effect is likely to be genuine and not due to reduced ability to infer crossovers within these regions, as the number of phase informative loci at these loci did not differ from the rest of the chromosome (Figure S14). It should be noted that in some chromosomes, female recombination rates dropped sharply in the first window of the chromosome (Figure 5), indicating that recombination rates are likely to be very low in close proximity to the centromere.

**Figure 4:**
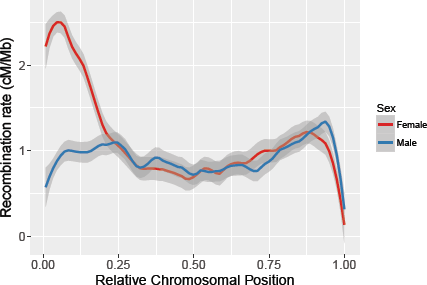
Loess smoothed splines of recombination rates across 32 acrocentric autosomes for males and females with a span parameter of 0.15. The centromere is assumed to be at the beginning of the chromosome. Splines for individual chromosomes are shown in Figure 5.

**Figure 5:**
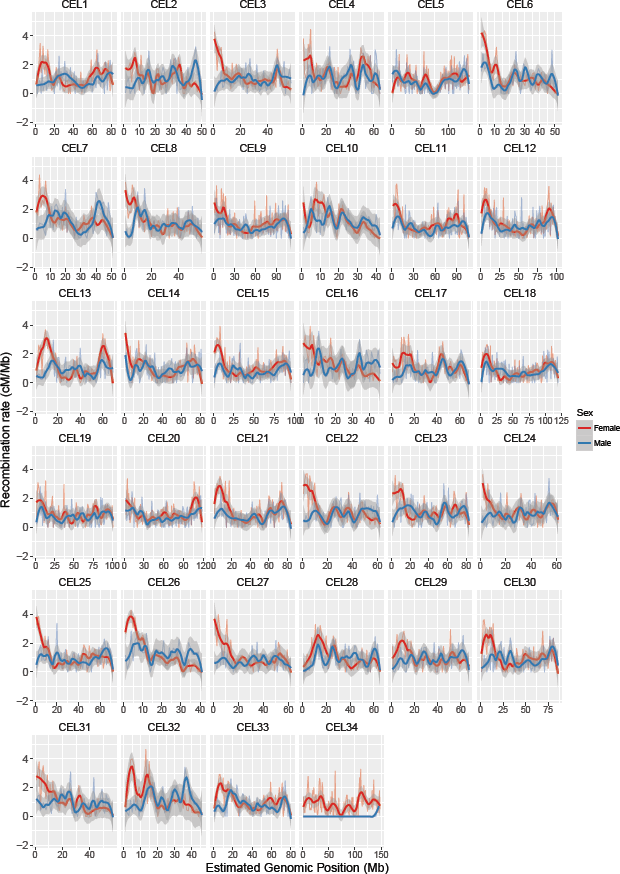
Loess smoothed splines of recombination rates in 1Mb windows across 33 autosomes for males and females with a span parameter of 0.2. All chromosomes are acrocentric with the centromere at the beginning of the chromosome (Gustavsson and Sundt, 1968), with the likely exception of CEL5. CEL34 is the X chromosome, with the pseudoautosomal region at the telomere end.

General additive models of recombination rate variation in acrocentric chromosomes indicated that female recombination rates at the closest proximity to the centromere were higher in fission chromosomes that would have had to have formed a new centromere (Figure 6A); this result held when one or two chromosomes were removed (not shown), and when considering small chromosomes only (Figure 6B). There were no differences in recombination rates in males with differences in chromosome history (Figure S15). There was no evidence of differences in transmission distortion with chromosome history in closest proximity to the centromeres in either sex, although there were subtle differences 5-6Mb from the centromere (P <0.05, Figure S16); full per-locus results are provided in Table S6.

**Figure 6:**
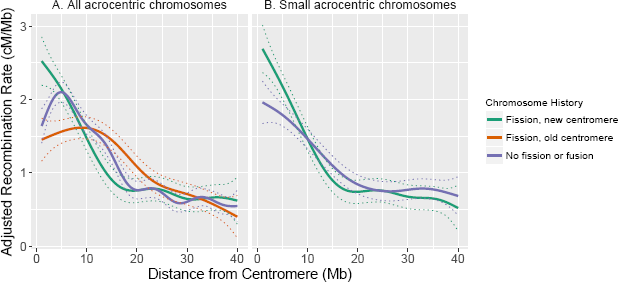
General additive model curves of adjusted recombination rate in females (k = 10). A. All acrocentric chromosomes, including fission chromosomes forming a new centromere (n = 6), fission chromosomes retaining the existing centromere (n = 6) and chromosomes with no fission or fusion (n = 20). B. Small acrocentric chromosomes, including fission chromosomes forming a new centromere (n = 5) and chromosomes with no fission or fusion (n = 6). Dashed lines indicate the standard errors. Recombination rates were adjusted for chromosome length (see main text).

## Discussion

In this study, we constructed predicted and skeleton linkage maps for a wild population of red deer, containing 38,083 and 10,835 SNPs, respectively. Females had higher recombination rates than males, which was driven by significantly higher recombination rates in pericentromeric regions. These rates were unusually high compared to other mammal species such as cattle, sheep and humans (Ma *et al.,* 2015; Johnston *et al.,* 2016; Kong *et al.,* 2010), and the effect was more pronounced in fission chromosomes that have formed centromeres more recently in their history. Here, we discuss issues related to the map assembly and utility, before proposing two explanations to explain strong heterochiasmy in peri-centromeric regions: (1) that this mechanism may have evolved to counteract centromeric drive associated with meiotic asymmetry in oocyte production; and/or (2) that sequence characteristics suppressing recombination in close proximity to the centromere may not have yet evolved at the neo-centromeres.

### Utility of the red deer linkage map

The final predicted linkage map included 38,083 SNPs, accounting for 98.8% of polymorphic SNPs within this population. Whilst several large-scale rearrangements were identified in the red deer lineage (Table 1), marker orders generally corresponded strongly to the cattle genome order. We are confident that the maps presented here are highly accurate for the purposes of genetic analyses outlined in the introduction; however, we also acknowledge that some errors are likely to be present. The limited number of meioses characterised means that we cannot guarantee a correct marker order on the predicted map at the same centiMorgan map positions, meaning that some small rearrangements may be undetected within the dataset. Furthermore, the use of the cattle genome to inform initial marker order may also introduce error in cases of local genome misassembly. Considering these issues, we recommend that the deer marker order is used to verify, rather than inform, any *de novo* sequence assembly in the red deer or related species.

### Mapping of the X chromosome (CEL34)

The X chromosome (CEL34) showed the highest level of rearrangement, including two translocations in the deer lineage, one of which was a small region in the pseudoautosomal region (PAR) remapped to the distal end of the chromosome (Figure S5). However, some caution should be exerted in interpreting whether these rearrangements relative to other species are genuine, as it has been acknowledged that the X chromosome assembly in cattle is of poorer quality in comparison to the autosomes (Zimin *et al.,* 2009; Ma *et al.,* 2015). The X chromosome showed a similar pattern to the autosomes in the relationship between estimated chromosome length (Mb) and linkage map length (cM, Figure 3). This may seem counter-intuitive, as recombination rates in the X should be lower due to it spending one-third of its time in males, where meiotic crossovers only occur on the PAR. However, female map lengths were generally longer, and 64% of the meioses used to inform sex-averaged maps occurred in females; furthermore, the female-specific map showed that the X conformed to the expected map length (Figure S12). Therefore, the linkage map length of the X is as expected; however, we acknowledge that some errors or inflation may be present on the X given that fewer informative meioses occur in non-PAR regions.

### Predicting centromere positioning on the deer linkage groups

Cytogenetic studies have shown that deer chromosomes are acrocentric (i.e. the centromere is situated at one end of the chromosome), with the exception of one unknown metacentric autosome, which is one of the physically largest (Gustavsson and Sundt, 1968). Our results suggest that the strongest candidate is CEL5, which has undergone a fusion event in the deer lineage (Table 1). Unlike other autosomes, this linkage group shows strong concordance between males and female cM maps (Figure 2), elevated male recombination rate at the chromosome ends and reduced recombination in a ~8Mb region that corresponds with the fusion site at the centromeric regions of BTA17 and BTA19 (Figure 5). On the acrocentric chromosomes, we have assumed that centromeres are at the beginning of each linkage group, based on synteny of centromere positions with the cattle genome (Ma *et al.,* 2015). There is evidence that centromeres can change position on mammalian chromosomes (Carbone *et al.,* 2006; Graphodatsky *et al.,* 2011). However, the frequency of this is sufficiently low, and recombination patterns so consistent in our dataset (Figure 5) that we believe our assumption is justified, particularly for chromosomes that have not undergone fission or fusion events in either lineage (Table 1). Six of the fission chromosomes (CEL6, CEL8, CEL16, CEL19, CEL22 and CEL26) would have had to form new centromeres in the deer lineage. Direct orientation with the cattle genome shows similar patterns of recombination to other chromosomes (Figure 5), indicating that telomeric regions have most likely not changed, and that centromeres have positioned themselves at the beginning of the chromosomes. Nevertheless, we acknowledge that confirmation of centromeric positions will require further investigation.

### Sexual dimorphism in recombination landscape: a consequence of centromeric drive?

Females had considerably higher recombination rates in peri-centromeric regions, resulting in female-biased recombination rates overall; recombination rates along the remainder of the chromosome were similar in both sexes, and lowest at the closest proximity to the telomere (Figure 4). Sampling identical numbers of males and females with replacement confirmed that this observation is unlikely to be a result of differences in sample sizes between the sexes (Figure S13). Identifying female-biased heterochiasmy is not unusual, as recombination rates in placental mammals are generally higher in females, particularly towards the centromere (Lenormand and Dutheil, 2005; Brandvain and Coop, 2012). Nevertheless, the patterns of recombination rate variation observed in this dataset are striking for several reasons. First, our findings are distinct from the other ruminants, namely cattle and sheep, which both exhibit male-biased heterochiasmy driven by elevated male recombination rates in sub-telomeric regions, with similar male and female recombination rates in peri-centromeric regions (Ma *et al.,* 2015; Johnston *et al.,* 2016). Indeed, all other mammal studies to date show increased male sub-telomeric recombination even if female recombination rates are higher overall (e.g. in humans and mice; Kong *et al.,* 2010; Liu *et al.,* 2014). Second, whilst female recombination rates tend to be relatively higher in peri-centromeric regions in many species, the degree of difference is relatively small compared to that observed in the deer, and is generally suppressed in very close proximity the centromere (Brandvain and Coop, 2012); in this study, we observed female recombination rates of 2-5 times that of males within this region.

There are several hypotheses proposed to explain increased recombination rates in female mammals (Brandvain and Coop, 2012). The most prevalent has been the idea that increased crossover number in females protects against aneuploidy (i.e. non-disjunction) after long periods of meiotic arrest during Prophase I (Morelli and Cohen, 2005; Coop and Przeworski, 2007; Nagaoka *et al.,* 2012). While we cannot rule this out as a potential driver, this hypothesis dominates the human literature, where females have one of the longest meiotic arrests of any mammal (Burt and Bell, 1987). Female deer on Rum reach sexual maturity at a relatively young age (~1.5 − 2.5 years of age) compared to other mammals, such as monkeys and great apes, and similar to other ruminants such as sheep and cattle (Burt and Bell, 1987). A more compelling hypothesis relates to the role of meiotic drive, where asymmetry in meiotic cell divisions in females can be exploited by selfish genetic elements associated with the centromere (Brandvain and Coop, 2012 & Introduction), where higher female recombination at the pericentromeric regions may counteract centromeric drive by increasing the uncertainty associated segregation into the egg (Haig and Grafen, 1991). In addition to a global mechanism driving local increases in recombination, our observation that chromosomes with newer centromeres show increased recombination in close proximity to the centromere may support this idea (Figure 6). Whilst there is still generally little consensus on the mechanisms related to the formation of new centromeres (Rocchi et al., 2011), conflict between centromeric proteins and repetitive centromeric DNA may lead to rapid evolution of centromere strength in a new or recently formed centromere (Rosin and Mellone, 2017). Increased recombination rates in close proximity to a newer centromere could provide a mechanism to counter stronger drive.

However, there are alternative (but not exclusive) arguments to this in the current dataset. The first is that increased recombination on new centromere chromosomes may be because sequence characteristics suppressing peri-centromeric recombination have not yet evolved in proximity to more recent centromeres. This is supported by our findings that there is no evidence of transmission distortion at centromeric regions on any of the chromosomes, although it also can be argued that centromeres have stabilised in the contemporary population, and that our current study cannot investigate historical differences in centromere strength. Additionally, the observed effect may be partially driven by mapping errors at the chromosome ends, particularly if polymorphisms in close proximity to the centromere have not been characterised and/or mapped.

### Conclusions and future directions

Our study has created a new linkage map resource for red deer and will facilitate genome-wide studies and genome assembly projects in red deer and related species. We have argued that increased recombination at peri-centromeric regions in females may be a mechanism to counteract meiotic drive; however, testing this hypothesis will require further investigation. Cytogenetic studies will allow confirmation of centromere positioning, and will give insight into how and why chromatin and cohesin structure does/does not suppress recombination in the pericentromeric region, and how their dynamics vary across the deer lineage (Vincenten *et al.,* 2015). In addition, *de novo* genome assemblies have the potential to verify map orders where possible; sequencing multiple deer genomes will allow us to determine population-scale recombination rates and hotspots (Chan *et al.,* 2012), with the potential to investigate historical variation in rate based on signatures of biased gene conversion (Capra *et al.,* 2013).

## Acknowledgements

We thank T. Clutton-Brock, F. Guinness, S. Albon, A. Morris, S. Morris, M. Baker and many others for collecting field data and DNA samples and their important contributions to the long-term Rum deer project. Discussions with C. Bérénos, J. Risse and M.D. Edge aided the study; we also appreciate discussion with G. Coop, T. Lenormand, L. Ross, D. Charlesworth, L. Ma and many Twitter users (Y. Brandvain, L. Holman, A. Kern, A. Mason, T. Price and L. Theodosiou) on interpreting the pattern of high peri-centromeric recombination in females. We thank Scottish Natural Heritage for permission to work on the Isle of Rum National Nature Reserve, and the Wellcome Trust Clinical Research Facility Genetics Core in Edinburgh for performing the genotyping. This work has made extensive use of the resources provided by the University of Edinburgh Compute and Data Facility (http://www.ecdf.ed.ac.uk/). The long-term project Rum deer is funded by the UK Natural Environment Research Council, and SNP genotyping was supported by a European Research Council Advanced Grant to J.M.P. S.E.J. is supported by a Royal Society University Research Fellowship.

## Author Contributions

J.M.P and J.H. organised the collection of samples. P.A.E. and J.H. conducted DNA sample 487 extraction and genotyping. S.E.J. designed the study, analysed the data and wrote the paper. 488 All authors contributed to revisions.

